# Physiological Dynamic Compression Regulates Central Energy Metabolism in Primary Human Chondrocytes

**DOI:** 10.1101/359885

**Authors:** Daniel Salinas, Brendan M. Mumey, Ronald K. June

**Keywords:** osteoarthritis, cartilage repair, mechanotransduction, chondrocyte, systems biology, metabolic flux analysis

## Abstract

Chondrocytes use the pathways of central metabolism to synthesize molecular building blocks and energy for cartilage homeostasis. An interesting feature of the *in vivo* chondrocyte environment is the cyclical loading generated in various activities (*e.g*. walking). However, it is unknown if central metabolism is altered by mechanical loading. We hypothesized that physiological dynamic compression alters central metabolism in chondrocytes to promote production of amino acid precursors for matrix synthesis. We measured the expression of central metabolites (*e.g*. glucose, its derivatives, and relevant co-factors) for primary human osteoarthritic chondrocytes in response to 0-30 minutes of compression. To analyze the data, we used principal components analysis and ANOVA simultaneous components analysis, as well as metabolic flux analysis. Compression induced metabolic responses consistent with our hypothesis. Additionally, these data show that chondrocyte samples from different patient donors exhibit different sensitivity to compression. Most important, we find that grade IV osteoarthritic chondrocytes are capable of synthesizing non-essential amino acids and precursors in response to mechanical loading. These results suggest that further advances in metabolic engineering of chondrocyte mechanotransduction may yield novel translational strategies for cartilage repair.

## 1 Introduction

Central energy metabolism is the primary set of pathways by which cells harvest energy and other molecular building blocks. These pathways include glycolysis, the pentose phosphate pathway, and the tricarboxylic acid cycle (Fig. 1). From these pathways, cells can generate both adenosine triphosphate (ATP), to fuel cellular processes, and precursors to non-essential amino acids. Therefore, these pathways are essential in the study of how chondrocytes regenerate articular cartilage. Note that we include respiratory pathways despite chondrocytes having been historically considered glycolytic cells. Recent data suggest that chondrocytes utilize both glycolytic and oxidative pathways to produce energetic particles and other useful precursors (Coleman et al 2016; Griffin et al 2012; Martin et al 2012; Zignego et al 2015; Blanco et al 2011).

**Fig. 1.**
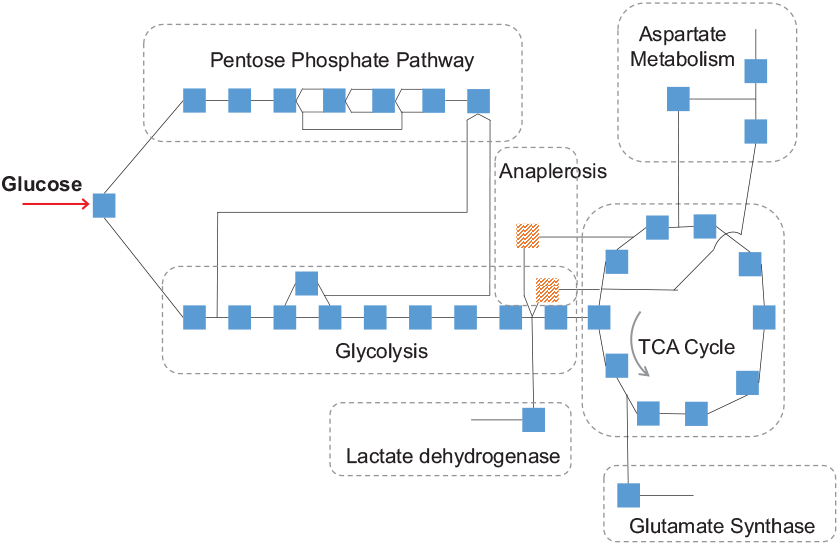
Map of central energy metabolism. Each square represents a modeled reaction, and lines represent metabolites, most of which are detected experimentally. Pathways included are glycolysis, pentose phosphate pathway, and the citric acid cycle (with anaplerotic reactions shown in red), and reactions in aspartate, glutamine, and lactate metabolism that share metabolites with these pathways. Electron transport chain reactions not shown for simplicity, but are included in the model.

The objective of this study was to analyze the metabolic changes induced in chondrocytes subject to physiological loading. Chondrocytes reside in a viscoelastic microenvironment (Alexopoulos et al 2005; Xu et al 2016). Everyday activities, such as walking, apply cyclical loads to chondrocytes, transmitted through the pericellular matrix (PCM). Because of its viscoelastic nature, the chondrocyte PCM is capable of both storing and dissipating the mechanical energy generated by *in vivo* activity. However, it is unknown how cyclical mechanical loading, in the short term and at physiological levels, affects central metabolism in chondrocytes.

We hypothesized that a short term physiological dynamic compression regime would alter central metabolism in chondrocytes to promote production of amino acid precursors. Recent studies (Coleman et al 2016; Brouillette et al 2014) link mechanical stimulation to central energy metabolism, demonstrating that injurious loading is inimical to respiration. Furthermore, studies in bovine and murine models have shown dynamic compression induces Ca^2+^ signaling (Wann et al 2012; Pingguan-Murphy et al 2006), which is a widely studied regulator of respiratory function in mammals (Griffiths and Rutter 2009). Our hypothesis that moderate, physiological stimulus promotes cartilage homeostasis is motivated by the larger question of whether moderate physical activity can be used to trigger tissue maintenance in diseased chondrocytes. Because the PCM is composed of proteins such as Type VI collagen and fibroblast growth factor family members (Wilusz et al 2014; Vincent et al 2007), we studied the production of amino acid precursors in central metabolism as indicative of the potential to maintain cartilage homeostasis.

To test this hypothesis, we analyzed metabolomics harvested from primary human chondrocytes obtained from osteoarthritic patients. To ensure accurate transmission of compressive loads, the chondrocytes were compressed while encapuslated in physiologically stiff agarose (Zignego et al 2015). Physiological cartilage stiffness ranges between 25 and 200 kPa, which can be achieved using 4.5% agarose gel. We analyzed the expression of 37 metabolites that participate in the central energy pathways listed above. We performed principal components analysis (PCA), ANOVA-simultaneous components analysis, and metabolic flux analysis with a recently developed stoichiometric model (Salinas et al 2017) to determine whether the changes observed over time were indicative of the production of amino acid precursors.

The results show that compression induces consistent changes in central energy metabolites after 30 minutes, and in some cases as little as 15. These results demonstrate that osteoarthritic chondrocytes can alter their central metabolism in response to applied compression. Furthermore, chondrocytes increase production of precursors to non-essential amino acids, suggesting a mechanistic link between compression and maintenance of cartilage homeostasis that involves central metabolism.

## 2 Methods

This study is an expanded analysis of data previously published. See (Zignego et al 2015) for additional details on methodology. These data come from primary human chondrocytes harvested from femoral heads of total hip arthroplasty patients under an IRB-exempt protocol.

### 2.1 Chondrocyte Encapsulation and Compression

The methodology for compressing chondrocytes has been previously published (Zignego et al 2014). Briefly, chondrocytes were harvested from five patients undergoing joint replacement surgery. All had stage IV osteoarthritis. Chondrocytes were isolated, expanded for one passage, and subsequently encapsulated in agarose gel of physiological stiffness (*~*45 kPa) to ensure transmission of compressive loads. After three days of equilibration in the gel, they were subjected to dynamic compression for up to 30 minutes. The compression was designed to simulate the human gait (1.1 Hz, *~*5% strain with 1.9% amplitude). Samples were collected at 0 (no compression), 15, and 30 minutes. Based on the expected PCM deposition (Dimicco et al 2007) and *in vivo* PCM stiffness (Darling et al 2010), we expect these strains to be in the physiological sub-injurious range (Chan et al 2016). We used independent samples from five donors, and had three to five replicates per donor.

### 2.2 Metabolite Extraction

The methodology for quantifying metabolites from central metabolism has been previously published (Jutila et al 2014). In summary, chondrocytes were flash-frozen after compression, and metabolites were extracted and quantified using high performance liquid chromatography coupled with mass spectrometry (HPLC-MS). A targeted analysis yielded intensities for 37 known metabolites in central metabolism.

### 2.3 Components Analysis

Metabolomic data describe metabolic state in terms of the abundance of each of its metabolites. We applied both principal components analysis (PCA) and ANOVA-Simultaneous Components Analysis (ASCA) to classify the ways in which metabolite abundances vary together. To apply PCA, the data were compiled into a matrix, **X**, with a row for each sample and a column for each metabolite. The dimensions of **X** in this study were 69 *×* 37, since 69 samples were harvested and each sample consisted of measurements for 37 metabolites. The entry at row *i* and column *j* is the abundance of metabolite *j* in sample *i* as measured by HPLC-MS. When two metabolite abundances vary together, this can be quantified using their covariance. The principal components are the eigenvectors of the covariance matrix of **X** after it has been mean-centered (Ten Berge et al 1992). The eigenvectors with the largest eigenvalues follow the largest covariation of abundances. The principal axes of variation are the eigenvectors with the largest eigenvalues.

To explain ASCA, we first define the ANOVA effects model. In this model, each value in **X** is interpreted as a sum of the effects of each factor in the experiment. In this study, the factors are the length of time the sample was compressed and the donor it was taken from. As a sum of effects, the abundance *x_ij_* of metabolite *j* in sample *i* is

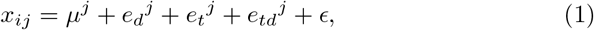

where *µ^j^* is the mean abundance of *j* over all samples, *e_d_^j^* is the effect the donor of sample *i* has on the abundance of *j*, *e_t_* is the effect compressing the sample for time *t* has on *j*, *e_td_* is the effect on *j* the interaction between donor and time (interpreted as donor sensitivity to compression), and *∈* is random error. The effects in eq. 1 are defined as follows,

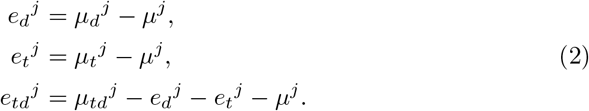

The mean *µ_d_^j^* is the mean of all samples from donor *d*, and the mean *µ_t_^j^* is the mean of the samples compressed for time *t*. Eq. 1 yields a set of matrices,

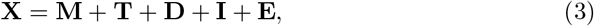

by computing the terms in eq. 1 for each *x_ij_* in **X** and arranging each collection of terms into a matrix according to **X**. There are several variations of ASCA (Timmerman and Kiers 2003), among them ASCA-P. We applied ASCA-P as in (Smilde et al 2005) by using the accompanying code, which applies PCA to each matrix on the right hand side of eq. 3. The data in each of the effect matrices varies only according to the effect. For example, the two terms *t_ij_* and *t_ko_* in **T** will differ only if sample *i* and sample *k* were compressed for different lengths of time. Thus, applying PCA to **T** captures how metabolites covary as a result of differences in compression time, but no other factors.

To illustrate PCA and ASCA, we consider the case where there are only three samples, all from a single donor, and two metabolites measured per sample. We let the data matrix be

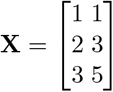

and let the time of compression for rows 1, 2, and 3 be 0, 15, and 30 minutes, respectively. Note that in this simple case, we assume no random error and have only time as a factor, resulting in **X** = **M** + **T**. The abundances of metabolites increases with compression time. The abundance of the first increases by 1 unit with each 15 minutes of compression, and the second increases by 2 units. The shift is therefore [0.1 0.2] over time, since multiplying this vector by the time yields the measured shift in abundance. The vector [0.1 0.2] is the direction along which the values of a sample in **X** change with time. The vector [0.1 0.2] is the first principal component as calculated by PCA, and it is also the first time component (first principal component of **T**) as calculated by ASCA. Both of these components match in this special case because mean centering **X** results in **T**.

### 2.4 Systems Analysis

In metabolic flux analysis (MFA), changes in abundance are interpreted as the result of reactions in metabolic pathways occurring at specific rates. The changes in abundance c can be computed from reaction rates **r** by

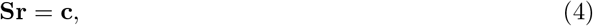

where **S** is the stoichiometric matrix of the network under study. A stoichiometric matrix is constructed from reaction stoichiometries, with a column for each reaction in the model and a row for each metabolite. For example, we will consider a system with only two reactions. Let reaction *r_i_* ∈ **r** be the first glycolytic reaction,

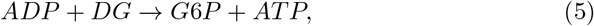

where *DG* corresponds to D-glucose and *G*6*P* corresponds to glucose-6-phosphate. Reaction *r_i_* therefore consumes 1 unit of *ADP* and one unit of *DG* and produces one unit of *G*6*P* and one unit of *AT P*. Let reaction *r_j_* ∈ **r** be the second glycolytic reaction

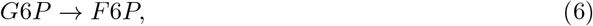

where one unit of *F*6*P*, fructose-6-phosphate, is produced from one unit of *G*6*P*. Assigning arbitrary rows and columns to metabolites and reactions, respectively, the stoichiometric matrix **S** is given below:

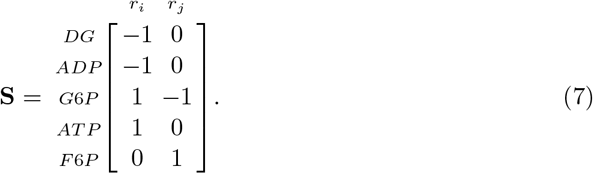

In this study, **S** was compiled from the reactions in glycolysis, the tricarboxylic acid cycle (with accompanying anaplerotic reactions), pentose phosphate pathway, and the electron transport chain. Due to the high abundance of lactate, glutamate, glutamine, aspartate and alanine in our samples, and their direct interaction with central metabolism, reactions mediated by lactate dehydrogenase, aspartate oxidase, aspartate ammonia lyase, glutamate synthase, and aspartate decarboxylase were also incorporated. The stoichiometric matrix used in our study can be found in the supplementary material (File S1).

Once the changes in abundance c have been computed, eq. 4 can be used to compute the rates that most closely match c while respecting thermodynamic constraints. The methodology used to solve for **r** is the same as our prior study (Salinas et al 2017). Briefly, we posed solving for **r** as a bounded variable least-squares (BVLS) problem. A BVLS problem is a least squares problem where variables can be constrained to be within lower or upper bounds. We used the bounded variable least squares (BVLS) solver included in MATLAB. The least-squares residuals were weighted by the inverse variance of their corresponding metabolite. Fluxes in r corresponding to irreversible reactions were constrained to have positive flux values.

We computed the changes in abundance over time **c** according to several criteria, solved for their respective **r**, and compared the results. We computed **c**:

1. using a naive approach: subtracting the median abundance at time 0 from the median at time 30, yielding a vector **c**_1_,
2. letting **c**_2_ be the third principal component resulting from PCA (the eigenvector of the covariance matrix with the third highest eigenvalue), and
3. letting c_3_ be the first principal component of the time factor matrix **T** using the effects model decomposition as implemented in ASCA.

We label the **r** computed from **c**_1_, **c**_2_, and **c**_3_ as **r**_1_, **r**_2_, and r_3_, respectively, and compare them in §3. Each axis of variation encodes a proportion of metabolites to each other. Interpreting PCA or ASCA axes as changes over time comes from how axes scores are computed. For example, assume sample **s_1_** has a higher score for axis **a** than sample **s_2_**, i.e. *s*_1_ is further along axis **a** than *s*_2_. This implies a larger inner product between **a** and **s_1_**. Given an axis **a**, any sample s_i_ can be expressed as a sum of an **a** component and a complement as follows:

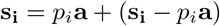

where *p_i_* is the inner product of **a** and **s_i_**. Thus, the difference **s_1_ *−* s_2_**, will have a positive a component:

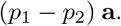

When samples that were compressed for a longer time have a higher *p_i_*, this implies that additional compression induced a change in the direction **a**. This implies that additional compression induced a chance in the direction **a**.

## 3 Results

### 3.1 Principal Components Analysis

PCA indicated that the variation between donors is larger than the variation that could be attributed to compression within each donor. It also suggested that, despite differing in base metabolite abundance, donors respond to compression in a similar manner (Fig. 2). Labeling the data according to donor and compression time showed that the axes of the first two principal components cluster points according to donor, with the third separating points within those clusters according to a mix of interdonor differences and compression time. The third PCA axis had loadings that showed negative coefficients for glucose, glutamate and aspartate, and positive coefficients for glutamine and pyruvate (Fig. 3A). Thus, samples that are further along the third PCA axis will have lower glutamate and aspartate, and higher glutamine and pyruvate, assuming they have equal scores on all the other PCA axes.

**Fig. 2.**
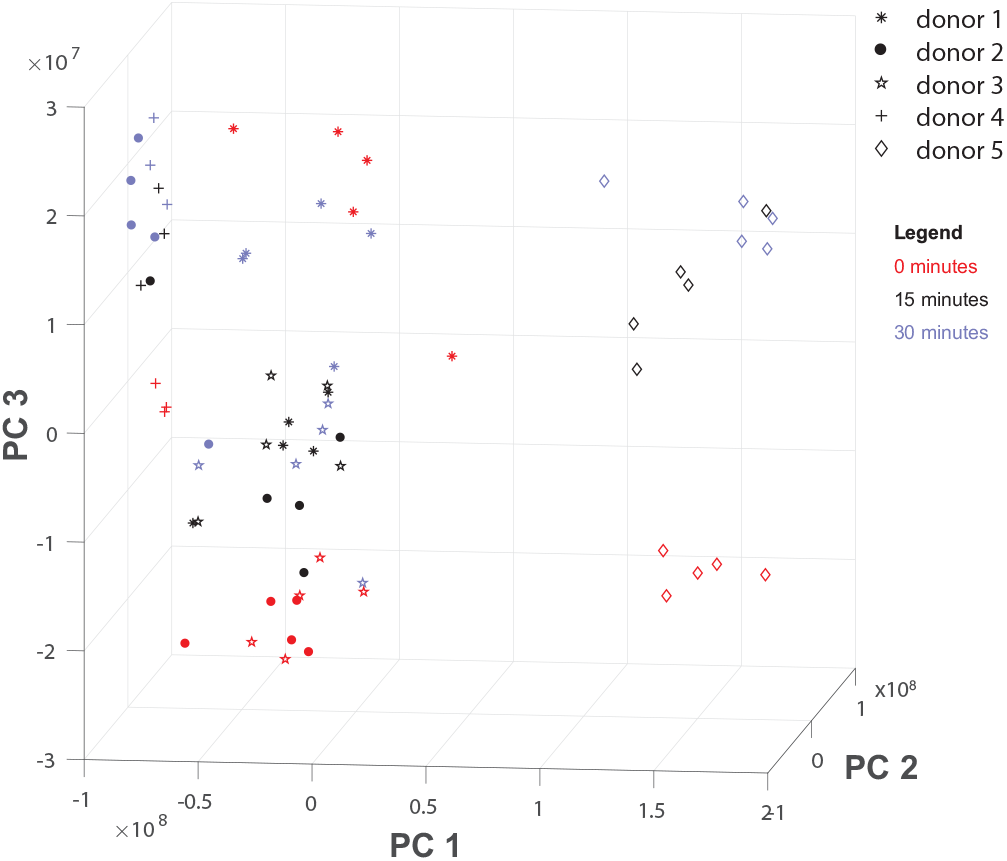
Principal components analysis of donor data. Each point represents a sample (associated with a donor and time of compression and composed of 37 measurements) after projection onto the three principal axes of variation. Red, black, and blue samples have been compressed for 0, 15, and 30 minutes, respectively. PC1 (88% variation explained) captures the differences between donor 5 and the rest of the donors; PC2 (9%) and PC3 (3%) capture the variation between donors and across compression time. PC3 especially is associated with the direction of change across time. Samples compressed for a longer time are higher along axis PC3 than those compressed for a lesser time, if they are from the same donor.

**Fig. 3.**
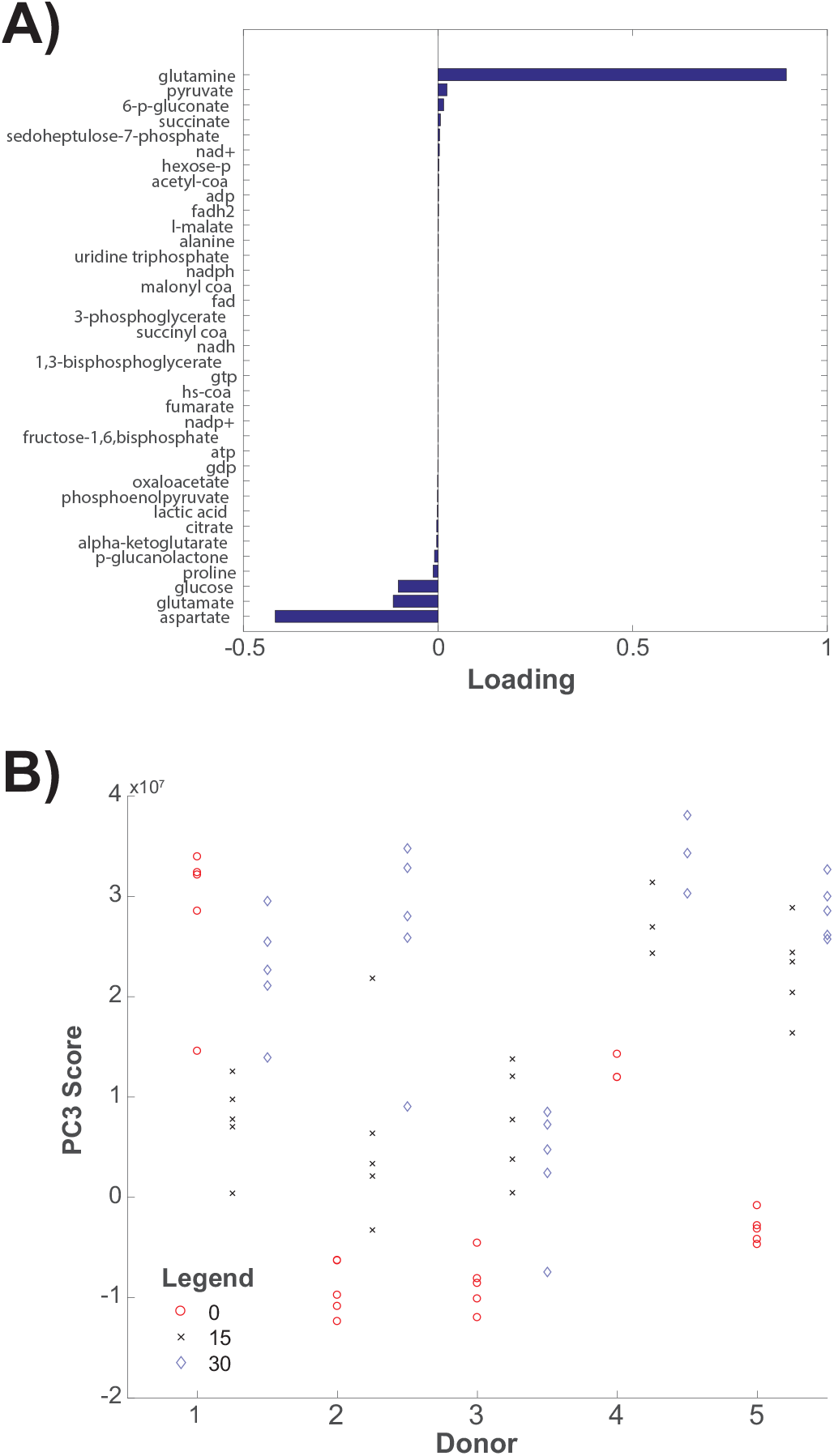
(A) Loading values along PC3 for each metabolite, sorted from least (bottom) to greatest (top). (B) Data for donors 1-5 after projection onto the third principal component. Red, black, and blue samples represent samples compressed for 0, 15, and 30 minutes, respectively. Compressed samples tend toward the top, obtaining higher PC3 scores.

Compressed samples tended to have a higher position along the third principal component axis than the samples that were not compressed, provided the comparison is between samples from the same donor (Figs. 2, 3B). The exceptions are donors 3 and 1. We explain these discrepancies in §4.

### 3.2 ANOVA Simultaneous Components Analysis

While the third axis of PCA is associated with changes induced by compression, we applied ASCA to rigorously define these changes according to the ANOVA model. ASCA showed the changes over time can be decomposed into two principal axes. The loadings for the first ASCA axis (which accounted for 89% of the time variation) shared the negative coefficients for glucose and positive coefficients for glutamine that the third PCA axis had (Fig 3A). However, the aspartate and glutamate coefficients are positive. Also, the coefficient for proline is the third largest. Despite these discrepancies, the axes have a high correlation (> 0.5) and can discriminate between non-compressed and compressed data (Figs. 4B, 5). However, the decline in score observed in donor 3’s samples is not present when plotted along the ASCA time axis, showing the ASCA axis to be a better discriminant. The difference in the loadings for glutamate and aspartate show that the PCA axis captured additional sources of variation. In particular, the axis captures the differences between donor 3 and donor 4 as well as those induced by compression time, as elaborated in §4.

**Fig. 4.**
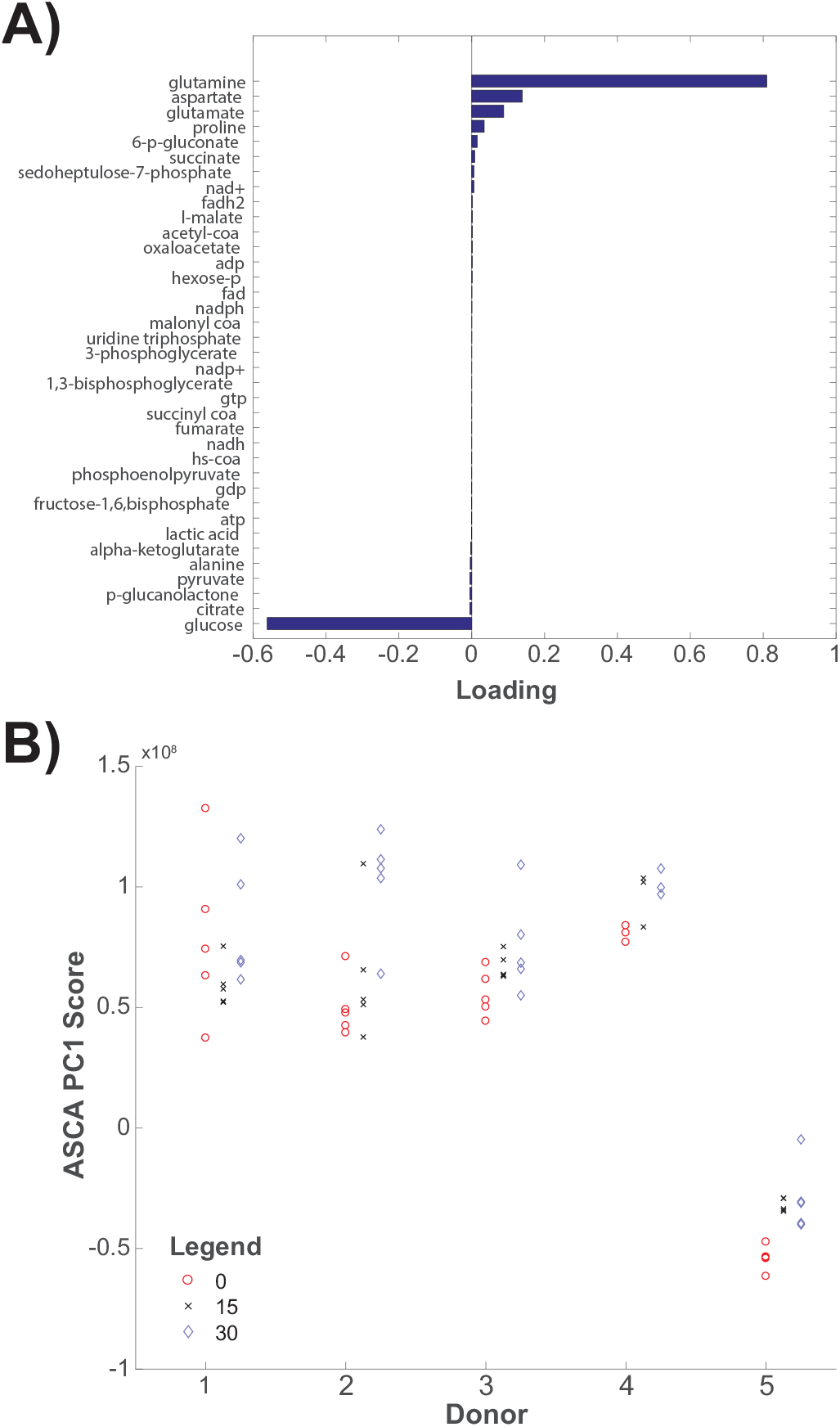
(A) Loading for each metabolite along the first principal component computed by ANOVA-Simultaneous Components Analysis (ASCA PC1), sorted from least (bottom) to greatest (top). (B) Data for donors 1-5 after projection onto ASCA PC1 (vertical dimension). Red, black, and blue samples represent samples compressed for 0, 15, and 30 minutes, respectively. Compressed samples tend toward the top, obtaining higher ASCA PC1 scores.

**Fig. 5.**
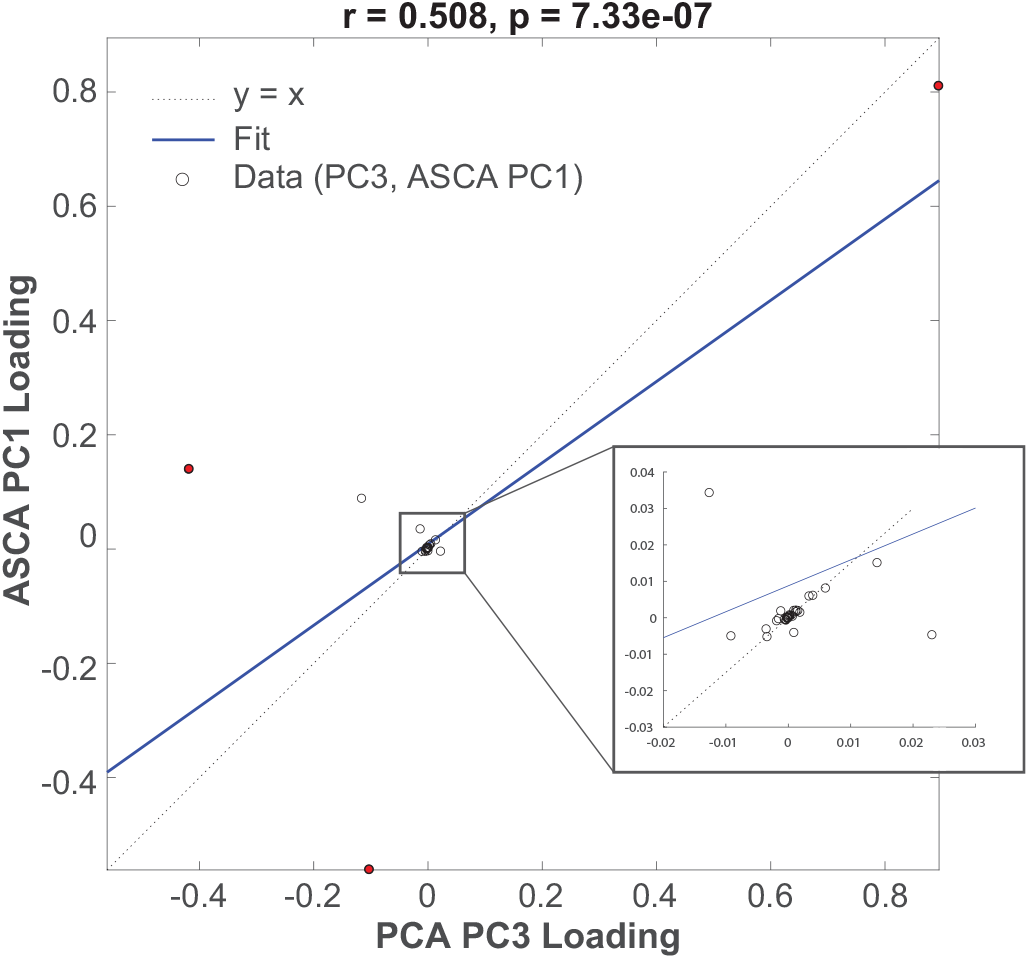
Regression analysis finds a relationship (r = 0.501, p¡0.001) between the third principal component (PCA PC3) and the first principal component of time (ASCA PC1). The correlation coefficient is 0.508 (*p <* 0.05). For each metabolite, the PC 3 loading is the horizontal coordinate and the ASCA PC 1 loading is the vertical coordinate. The main discrepencies relate to metabolites with either heavy production (aspartate and glutamine) or consumption (glucose). These are plotted in red.

The significance of the difference in scores along the time axis was assessed via a label permutation study as in (Vis et al 2007). Briefly, the permutation study measures significance by studying the difference in group variance. By reassigning the samples to different groups (i.e. changing a sample’s donor or compression time), a distribution of the variance of each group is formed over all possible label permutations. If samples belong to different populations, the variance within groups will be smaller than the vast majority of the variances observed when assigning the samples to different groups. The permutation study showed that the variation of compressed and uncompressed groups from the mean was much larger than could be expected by random variation due to small sample size, to high significance (*p* = 0.00074). This indicates that both donor and compression time affect chondrocyte central metabolism.

### 3.3 Metabolic Flux Analysis

As discussed in §2, the changes **c**_1_ in metabolite abundances allow us to infer the reaction rates **r**_1_ that produced them using MFA. The change in abundances over time corresponding to the first ASCA axis requires high activities for glycolysis and the pentose phosphate pathway coupled with high conversion of ammonia lyase into fumarate, high conversion of malate to pyruvate, and high oxidation of aspartate (Fig. 6A). The rates that match the third PCA axis were almost identical to the ASCA rates (not shown). Rates 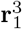, 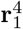, and 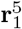 calculated from the median samples for donors 3, 4, and 5, respectively, were consistent with the ASCA rates, showing that ASCA infers the change in overall metabolomic profiles (Fig. 6B). The rates have two major clusters: one for donors 1 and 2 and the other for the donors 3, 4, and 5 (Supplement S2). Donors 1 and 2 are the youngest and oldest donors, respectively. Further discussion of these rates is found in §4.

**Fig. 6.**
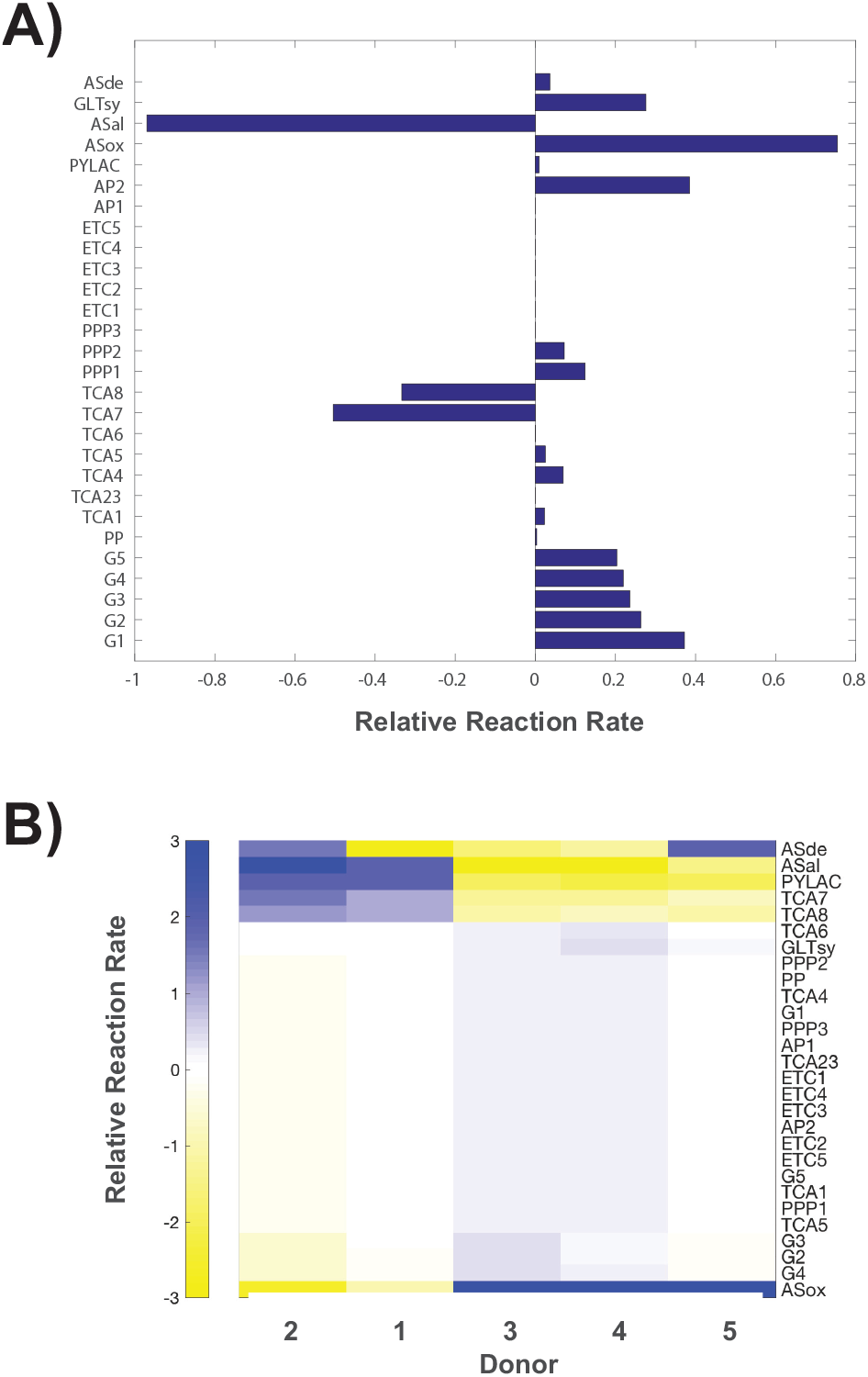
(A) Reaction rates that approximate the changes in metabolite abundances induced by 30 minutes of compression as computed by ASCA, **r**_3_. These rates are consistent with increased synthesis of aspartate and glutamine, all needed for protein synthesis as required by chondrocyte homeostasis. (B) Reaction rates 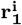 for each donor *i*, for the periods of 0-30 minutes of compression, calculated from the differences in the medians of the samples for each time point. These are consistent with the compression rates, demonstrating that the overall shift in metabolism within donors comes as a result of compression over time. Reaction labels defined in Supplement S1.

## 4 Discussion

The effect of physiological compression on the central metabolism of primary osteoarthritic chondrocytes was quantified using ASCA on targeted metabolomic data and interpreted via MFA. While inter-donor differences were larger than the differences in metabolism induced by compression, all donors reacted similarly to compression. The significance of this data is the demonstration that the central energy metabolism of human osteoarthritic chondrocytes responds to physiological compression. In this section, we discuss whether the data is consistent with protein precursor synthesis and the potential for load-induced cartilage repair.

### 4.1 Healthy Chondrocyte Response to Compression

For reference, we investigated the response of healthy chondrocytes to dynamic compression in the literature. We identified two studies that subjected primary human chondrocytes to dynamic compression regimes comparable to those in our study (Demarteau et al 2003; Grogan et al 2012). Because in (Grogan et al 2012) the compressive regime is supplemented with perfusion, we focus on the effects of compression as reported in (Demarteau et al 2003). A compression regime with 5% strain and 5% amplitude at 0.1Hz was used, comparable to the 5% with 1.9% amplitude and 1.1Hz used in our study. However, the total duration of the compression was substantially longer. Their study compressed chondrocytes for two hours at a time, in contrast to the 30 minute maximum used in our study. In Demarteau et al (2003), the question of whether chondrocyte compression induces the synthesis and accumulation of GAG was investigated. Both synthesis and accumulation were found to be dependent on the presence of GAG prior to loading. GAG synthesis was greater in chondrocytes that were compressed than in free-swelling controls once enough GAG had been synthesized by the acclimating chonrdocytes during the culture period. The measured response of increased GAG synthesis is in consensus with (Grogan et al 2012) and even various bovine studies, e.g. (Buschmann et al 1995). In these studies, GAG synthesis is assumed to be indicative of aggrecan anabolism. As indicated in (Esko et al 2009), aggrecan is the major proteoglycan in cartilage, and therefore GAG synthesis is meant to be interpeted as cartilage renovation.

### 4.2 Components Analysis

ASCA identified glutamine, aspartate glutamate, and proline as having a positive association with compression time, and it identified glucose as having a negative covariance. While donors have different ranges of scores, samples from a given donor that have been compressed for 30 minutes receive consistently higher scores than the uncompressed samples from the same donor. Donor 1’s uncompressed samples require further discussion.

Donor 1, the youngest and only male donor had uncompressed samples that received higher scores than even those samples that were compressed for 30 minutes. However, his uncompressed samples had the widest range, receiving both the lowest and highest scores of all donor 1’s samples. The 15 minute samples have a much, and on average receive lower scores than the samples compressed for 30 minutes. This suggests that, while the uncompressed samples were heterogeneous, compression induced homogeneous behavior, and this behavior was consistent with the behavior observed in the older donors’ samples. We observe that aging decreases aggrecan synthesis rates (Verbruggen et al 2000), and that impaired synthesis is one of the pathologies linked to OA incidence (Martin and Buckwalter 2002). Thus, it is possible that a wide range in ASCA 1 scores is indicative of the presence of chondrocytes in various stages of synthetic activity due to differences in senescence. We note that dynamic compression may synchronize chondrocyte metabolism, causing the subsequent homogeneity at 15 and minutes of compression. However, further analysis is required to determine if the observed heterogeneity stems from age or gender based effects.

Although the third PCA axis and the first ASCA time axis both correlate with time of compression, their discrepancy arises from the inclusion of donor variation when applying PCA. Specifically, aspartate, glutamate, and glucose deviate from the observed correlation (Fig. 5). This likely reflects their strong negative PCA axis 3 loading values (Fig. 3A). Importantly, the PCA axis 3 loading values are not independent of donor. Therefore, it is likely that these outliers reflect a combination of donor-dependent effects and large PCA axis 3 loadings.

We verified this by sorting the donors according to average score on PCA axis 3. On average, the lowest scoring donor, donor 3, has a higher concentration of glutamate and aspartate than the the highest scoring donor, donor 4. A loss of aspartate and glutamate results in negative loadings for glutamate and aspartate for the third PCA axis, in contrast to the positive loadings observed in the first ASCA time axis. However, on average, the samples compressed for 30 minutes had higher glutamate and aspartate abundances than the uncompressed samples, thus verifying that the first ASCA time accurately captures the variation over time. The correlation of the third PCA axis to the time of compression stems from the changes in the abundance of other metabolites (Figure 5), rendering it a mixture of the time and donor variations.

### 4.3 Metabolic Flux Analysis

Analysis of the reaction rates computed via MFA suggest a mechanism of glutamine and aspartate synthesis. We computed rates **r**_3_ that approximate the changes observed over compression time as defined by the first ASCA axis (Figure 6A). Glycolysis is prominent in this profile, resulting in ATP and pyruvate synthesis (Figure 7). Aspartate is synthesized from fumarate, which is synthesized from malate. TCA cycle malate is restored via pyruvate (AP2), indicating that overall the cells generate aspartate from a fumarate precursor. Also, fumarate is not generated via TCA cycle intermediates, rather through the action of the malic enzyme. Glutamine is synthesized from glutamate through the action of glutamate synthetase, and is aided by the malic enzyme in the process. Additionally, glutamate abundance also increases over time, though at a much lower rate, suggesting that synthesis from glutamate synthetase is an incomplete explanation. Glutamine may be an essential amino acid to chondrocytes (Handley et al 1980); the media contained glutamine, suggesting that the glutamine increase may also be due to transportation into the chrondrocytes rather than synthesis from glutamate. Glutamate synthesis may also be occurring due to reactions outside our model. Glutamate is a precursor to glutathione, and chondrocytes are known to undergo oxidative stress when compressed (Brouillette et al 2014). Aspartate and glutamine are precursors, and an increase in their synthesis may be tied to the production of the core protein in aggrecan. The core protein requires asparagine to bind N-linked oligosaccharides Vertel (1995). Whether aggrecan or not, an increase in protein synthesis is consistent with our previous study conducted on SW1353 chondrosarcoma (Salinas et al 2017). These data also show that compression induces a higher abundance of glutamine in the agarose-encapsulated chondrocyte samples.

**Fig. 7.**
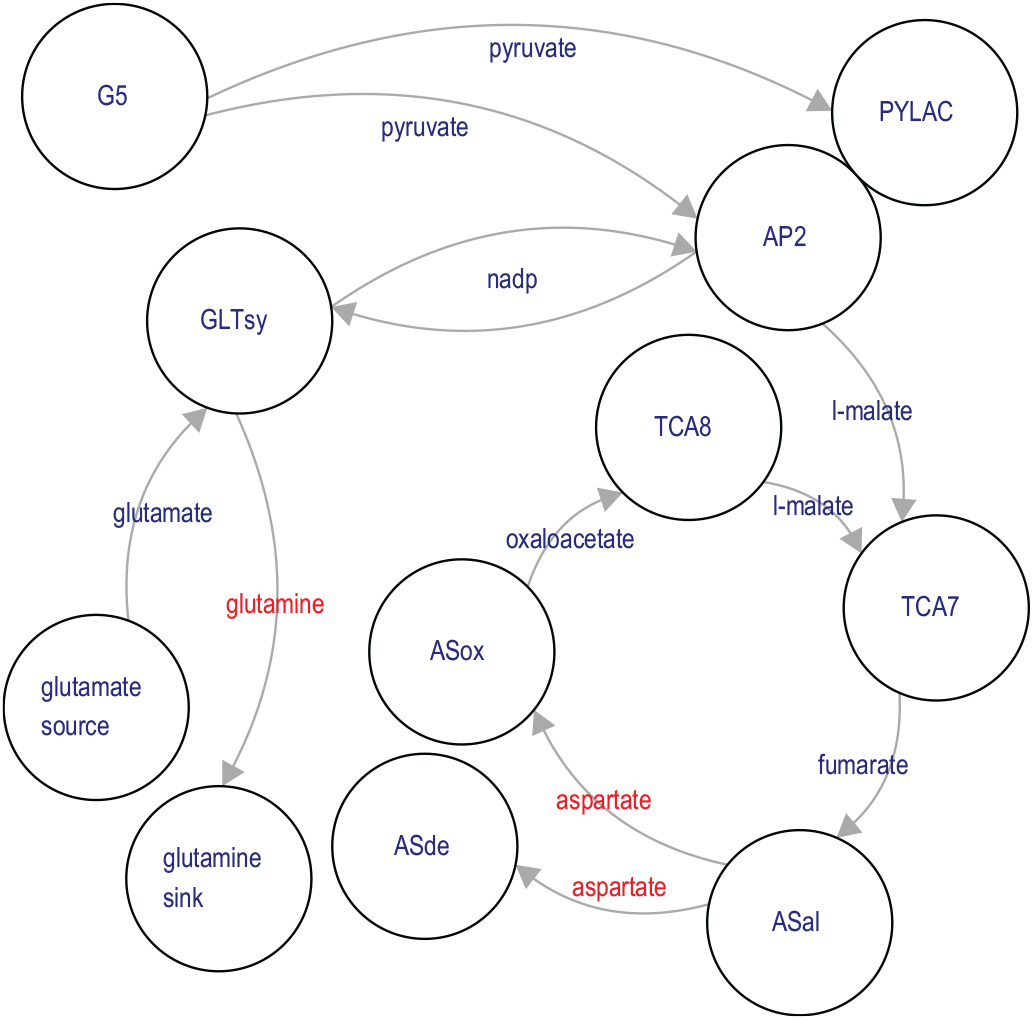
Flow of metabolites in compression-induced metabolic production of aspartate and glutamine. Network synthesis of aspartate and glutamine is accomplished via glutamate synthase (GLTsy) and the tandem action of ASal (adenylsuccinate lyase and adenylosuccinate synthetase isozyme 1). Synthesis of glutamine is aided by the action of the malic enzyme (AP2), which recycles the NADP^+^ produced by GLTsy into the NADPH required by GLTsy.

We computed reaction rates 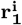 for central energy metabolism from each donor’s observed metabolic profile (Figure 6). The reaction rates computed from the median metabolite values from donors 1 and 2 cluster independently, indicating that within-donor variations were greater than compression-induced variations for these two donors. We computed rates for both the 0-15 and 15-30 periods of the experiment. However, due to the marked differences of cells from these donors between the 0-15 and 15-30 minutes of compression, we chose to compare the uncompressed control cells to cells compressed for 30 minutes for data reported in this manuscript.

## 5 Conclusion

These results demonstrate that even grade IV osteoarthritic chondrocytes are able to respond to physiological levels of compression by synthesizing non-essential amino acids. Future studies may build on these results by comparing these data to age- and gender-matched chondrocytes from healthy donors to use metabolic engineering to harness *in vivo* loading for cartilage repair. Future studies may also focus on maximizing the compression-induced production of metabolic precursors to amino acids found in this study. These analyses demonstrate the importance of using systems approaches to understand metabolomics data, as the integration of the data with these models can yield insight that is not clear from examining metabolite values in the absence of such a stoichiometric model.

## 6 Acknowledgements

This study was funded by the National Science Foundation (1342420, 1554708 and 1542262) and the NIH (P20GM103474).

## S1

Stoichiometric matrix used for metabolic flux analysis and flux balance analysis. An explanation of the reaction abbreviations used in the paper is included.

## S2

Values for reaction fluxes over the first and second 15 minutes of compression for each donor. Fluxes that generate the third PCA axis and fluxes that generate the first ASCA axis are included.

## S3

Values for PCA and ASCA decompositions of variation. The largest three components are included for PCA, and the largest two for ASCA.

